# Validation of a Miniaturized Permeability Assay Compatible with CRISPR-Mediated Genome-Wide Screen

**DOI:** 10.1101/471854

**Authors:** Claire Simonneau, Junning Yang, Xianguo Kong, Robert Kilker, Leonard Edelstein, Paolo Fortina, Eric Londin, Arie Horowitz

## Abstract

The impermeability of the luminal endothelial cell monolayer is crucial for the normal performance of the vascular and lymphatic systems. A key to this function is the integrity of the monolayer’s intercellular junctions. The known repertoire of junction-regulating genes is incomplete. Current permeability assays are incompatible with high-throughput genome-wide screens that could identify these genes. To overcome these limitations, we designed a new permeability assay that consists of cell monolayers grown on ∼150 μm microcarriers. Each microcarrier functions as a miniature individual assay of permeability (MAP). We demonstrate that false-positive results can be minimized, and that MAP sensitivity to thrombin-induced increase in monolayer permeability is similar to the sensitivity of the measurement of impedance. We validate the assay by showing that the expression of single guide RNAs (sgRNAs) that target genes encoding known thrombin signaling proteins blocks effectively thrombin-induced junction disassembly, and that MAPs carrying such cells can be separated effectively by a fluorescent probe from those that carry cells expressing non-targeting sgRNAs. These results indicate that MAPs are suitable for high-throughput experimentation and for genome-wide screens for genes that mediate the disruptive effect of thrombin on endothelial cell junctions.

## INTRODUCTION

One of the new opportunities afforded by the sequencing of the human genome, as well as of many others, is the ability to run genome-wide reverse screens to identify genes that underlie diverse phenotypes. The repurposing of CRISPR/deactivated (d) Cas9 for targeting either repressive or activating modules to genomic loci provides a more versatile and specific tool than RNAi, becoming the screen method of choice^1^. In the few short years since this tool became available, it has already been applied to numerous ends, including cancer drug discovery^2^, analysis of unfolded protein response^3^, and the identification of cancer immunotherapy targets^4^. CRISPR/dCas9-mediated screens take advantage of the accurate sgRNA-dependent targeting of protein modules that either repress or activate transcription to proximal upstream and downstream regions near the transcription start site. Transcriptional activation has been achieved by various combinations of a range of modules, including NF-κB trans-activating subunit p65^5^, tetrameric VP16 transactivation domain (VP64), heat shock factor 1, the catalytic histone acetyltransferase core domain of the human E1A-associated protein p300, and the viral replication and transcription activator [reviewed in^6^]. In contrast, transcriptional repression has been produced by a single module, the Krüppel-associated box (KRAB) domain of the zinc-finger transcription factor Kox^7^. The efficacy of the regulatory dCas9-fused module is highly dependent on the selection of the sgRNA genomic targets relative to the transcription start sites, based on rules that were derived and optimized empirically^5,8^. Activation on the order of 10^3^-fold, as measured by quantifying mRNA levels of selected genes, was achieved by some of the above complexes^6^, whereas transcriptional repression measured in the same way was on the order of 10^2^-fold, and rarely 10^3^-fold^9^.

Genome-wide screens employing activating or repressive dCas9 constructs and sgRNA libraries were deployed to identify toxin or drug resistance and vulnerability-conferring genes^9–14^, genes that mediate cell immune and inflammatory responses^15–18^, and genes required for cell survival, proliferation, and metastasis^10,19–22^. The majority of screens performed to date phenotyped cell enrichment or depletion, though some studies measured the rates of differentiation^23^, cell motility^24^, and the abundance of fluorescent protein^25^. In all cases, the quantified phenotypes were individual-cell attributes. Such assays are suitable for the pooled format that, with rare exceptions^26,27^, characterizes the vast majority of currently available sgRNA libraries, and enables a relatively facile and economical quantification of phenotypes. Though effective and fruitful, the current limitation to individual-cell functional assays leaves out collective cell functions which underlie physiologically essential processes. Here we validate a new functional assay we designed to facilitate adaptation of the CRISPR-mediated screen for the identification of genes that regulate the permeability of endothelial cell (EC) monolayers. The integrity of this monolayer is critical to the normal function of the vascular system because it confers the barrier function of the vessels. Since administration of thrombin produces large gaps in EC monolayers that can be detected with relative ease, we chose to validate the assay by testing if sgRNAs from an existing repressive CRISPR library^8^ designed to knockdown the transcription of thrombin signaling pathway genes, become enriched in unresponsive cells. Thrombin-induced permeability is not manifest in vertebrates under normal conditions, but does occur under trauma, sepsis, and other inflammation-inducing conditions in both animal models^28–30^ and patients^31,32^.

Currently available permeability assays are not compatible with the requirement of genome-wide screens because they cannot accommodate the large number of samples, and because their multi-well format is unsuitable for the pooled sgRNA libraries used in practically all screens. To our knowledge, the highest sampling capacity currently available for the measurement of paracellular permeability is provided by the 96-well plate filter insert assay format. A comprehensive screen of the human genome by 10 sgRNAs and 5 technical replicates per gene using this assay would require approximately 10^4^ multi-well plates, a number that is beyond the capacity of most research laboratories. To design a more feasible assay, we took advantage of the microcarrier-based cell culture technique developed to maximize the number of adherent cells grown per volume of medium. Typically, microcarriers consist of spherical beads of a diameter in the range of 100-200 μm, made of cross-linked polymers. Commercially available microcarriers come in a variety of polymeric matrices and surface treatments or coatings^33^. Microcarriers carrying EC monolayers had been used to study the effect of various agonists on cell monolayer permeability by measuring the absorbance of dye into microcarriers, or in a chromatographic format by measuring the elution time of a dye^34–37^

## RESULTS

### EC monolayers grown on single microcarriers serve as individual permeability assays

We sought an in vitro permeability assay that would be compatible with the high throughput format required for genome-wide screens to interrogate signaling pathways that regulate the integrity of endothelial cell junctions. In the new assay, each microcarrier is treated as an individual MAP. This facilitates high throughput testing of a large number of samples in a relatively small volume of growth medium (approximately 6.2×10^5^ microcarriers per 100 mL). We envisioned that ECs will be transduced by repressive (CRISPRi) sgRNA libraries and grown as clonal populations on each microcarrier. When treated by agonists that disrupt cell junctions, e.g. vascular endothelial growth factor (VEGF) or thrombin, junctions among ECs expressing sgRNAs that target genes required the for the cellular response to the administered agonist would fail to disassemble. To distinguish between microcarriers carrying responsive or non-responsive EC monolayers, we selected a probe that would bind to the microcarrier surface exposed through the gaps between responsive cells. Since the microcarrier type that supported EC growth optimally is coated with gelatin, we selected the collagen-binding fluorescently-conjugated fragment of fibronectin (FN_c_)^38^ as a probe because of its high binding affinity to gelatin^39^. The fluorescence of MAPs carrying responsive EC monolayers would increase upon thrombin treatment because of the binding of fluorescently-conjugated FN_c_ (FN_cf_) to the newly formed gaps between ECs. The darker MAPs that carry non-responsive monolayers would then be separated from the brighter ones by fluorescence-assisted sorting (Fig. 1).

**Figure 1:**
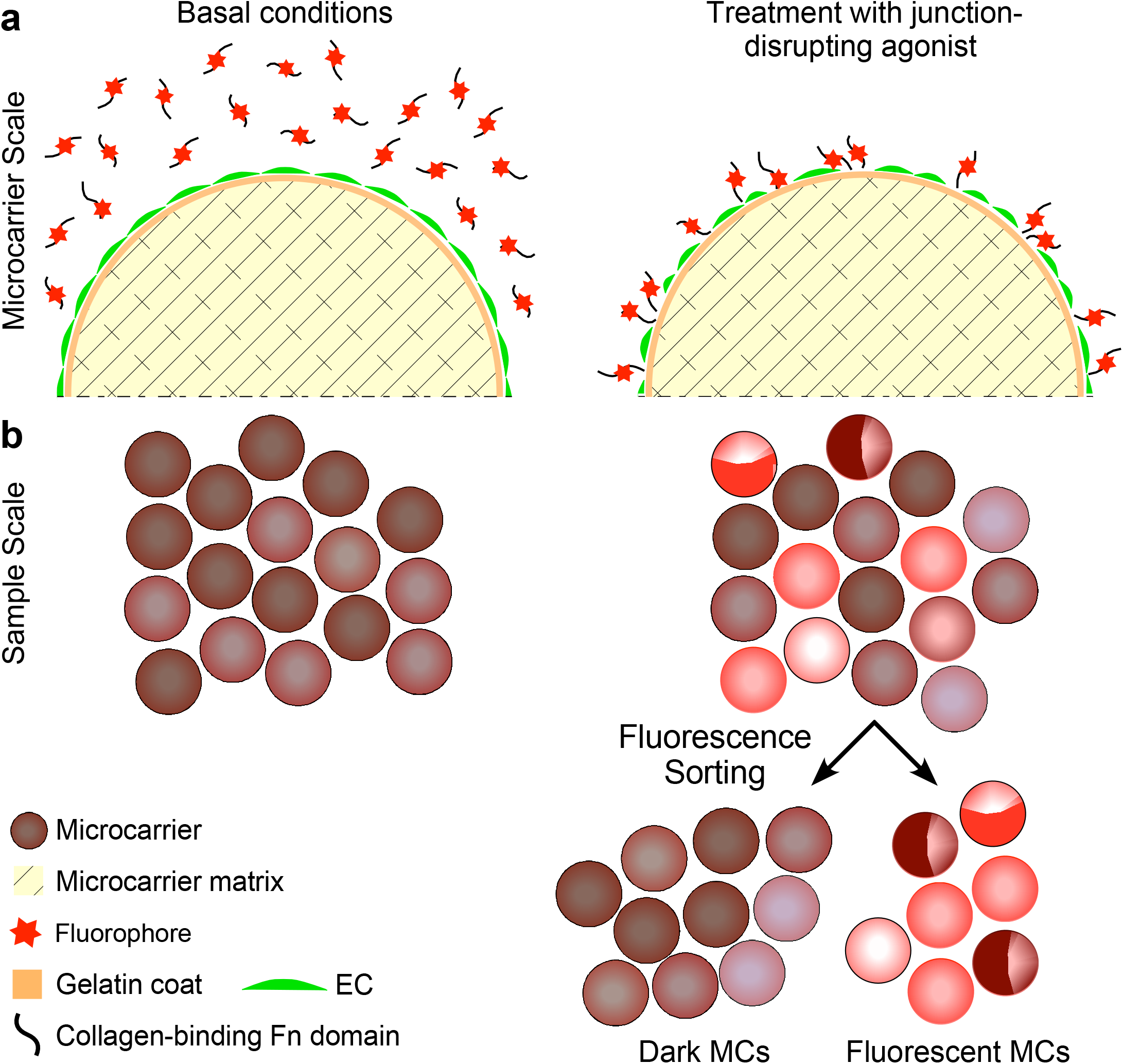
Microcarriers are repurposed as miniature individual permeability assays. **a.** Gelatin-coated microcarriers composed of cross-linked dextran carry a confluent EC monolayer (green, to designate calcein-loaded ECs), incubated in medium containing the collagen-binding proteolytic fragment of fibronectin conjugated to a fluorophore (FN_cf_). Once treated by a junction-disrupting agonist (thrombin in this study), FN_cf_ binds to the exposed gelatin surface between cells whose junctions disassembled in response to the agonist. **b.** Microcarriers carrying untreated ECs bind a minimal amount of FN_cf_. Microcarriers carrying agonist-treated ECs bind varying amounts of FN_cf_, depending on the identity of the sgRNA expressed by the clonal cell population on each microcarrier. Microcarriers carrying ECs that express sgRNAs targeting genes that encode proteins required for the induction of the disassembly of cell-cell junctions bind a low amount FN_cf_, similar to untreated microcarriers. ECs expressing sgRNAs that are unrelated to the signaling pathway of the junction-disrupting agonist respond by disassembling their junctions. FN_cf_ binds to the gelatin surface exposed between the responsive ECs, rendering the microcarriers that carry these cells fluorescent. The fluorescent microcarriers and the dark microcarriers are separated by fluorescence-assisted sorting. The gates of the sorting machine can be set up to capture any group of interest in this population, based on microcarrier fluorescence intensity.

### Thrombin increases the permeability of telomerase-immortalized human primary EC monolayers

Among a variety of known permeability factors, thrombin induces relatively large openings between confluent ECs^40^, on the same type of microcarriers used in this study^35^. To gauge thrombin’s effects on monolayers of telomerase-immortalized (human dermal) microvascular endothelial (TIME) cells, we measured its impact on permeability by two approaches. In the first, we measured the penetration of a fluorescently-conjugated dextran probe through an EC monolayer. In the second, we measured thrombin-induced change in monolayer impedance. Both approaches yielded large changes in the measured attributes, indicating substantial increases in monolayer permeability. In the first type of assay, the fluorescence of the probe in the lower chamber of the assay almost plateaued after a more than a 3-fold increase (Fig. 2a). In impedance measurements, thrombin concentrations of 0.5, 1.0, and 2.0 U/mL reduced the impedance by approximately one unit, although their rate of recovery to the pre-thrombin level became slower when thrombin concentration was increased (Fig. 2b). The impedances recovered their pre-thrombin levels and stabilized in approximately 120 min. To visualize the structural effects of thrombin on the EC monolayer, we immunolabeled the cells to track the localization of vascular endothelial cadherin (VEcad) and used phalloidin to observe thrombin-induced changes in the actin cytoskeleton. Thrombin induced opening of large gaps, which reached in some locations a width of approximately 10 μm, and formation of denser and thicker stress fibers (Fig. 2c), similar to previous observations^41^.

**Figure 2:**
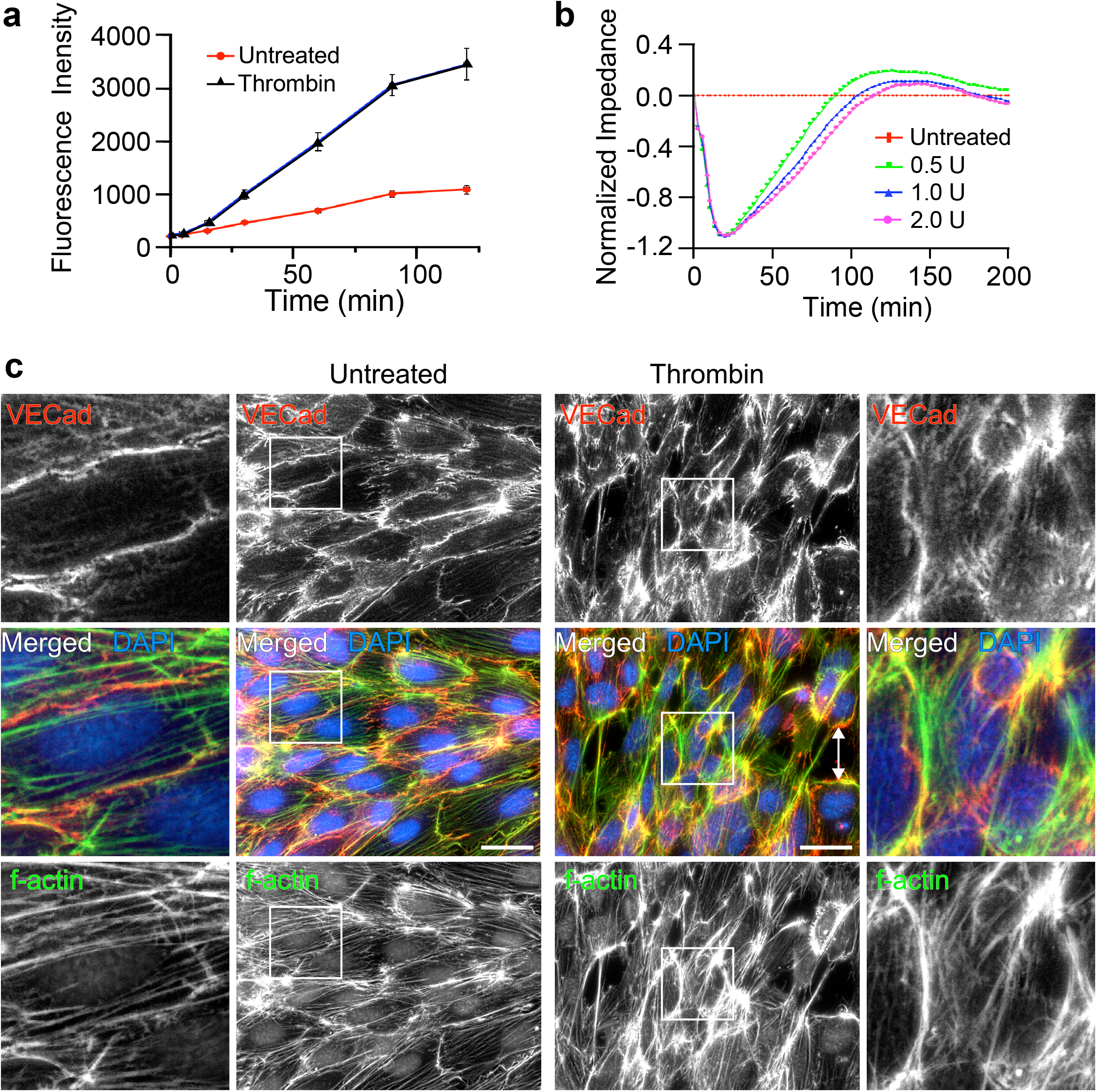
Thrombin increases the paracellular permeability of TIME cell monolayers. **a.** The permeability of TIME cell confluent monolayers grown on filter inserts to 2000 kDa FITC-dextran increased in response to treatment by 2.0 U thrombin applied at t=0 min for the duration of the experiment, as indicated by the augmentation of the fluorescence signal from the bottom well under. **b.** The impedance of TIME cell monolayers grown on electronic 16-well plates fell sharply after treatment by the indicated dosages of thrombin (applied as above), indicating that the integrity of the monolayer deteriorated. **c.** Epifluorescence images of confluent TIME cells in monolayers grown on gelatin-coated coverslips. Intercellular junctions disassembled upon treatment by 2 U/mL thrombin (Thr), as indicated by the change in VEcad signal. The cells formed thicker stress fibers and detached from each other. Magnified images of the regions marked by white frames are shown on the left and right sides. The double-pointed arrow marks a gap between thrombin-treated cells. Scale bars, 50 μm.

### Single TIME cells grow to confluence on gelatin-coated microcarriers

Following tests with several types of microcarrier polymers (e.g. polystyrene, diethylaminoethyl dextran), and coatings (e.g. fibronectin, collagen I), we determined that gelatin-coated microcarriers consisting of cross-linked dextran matrix are optimal for TIME cell growth. To observe monolayer integrity, we stained them with calcein, visualizing both the cell body and intercellular junctions (Fig. 3a). All the cells grown on each microcarrier in a genome-wide screen should express a single sgRNA to avoid mixed effects of silencing the expression of potentially antagonistic genes or additive effects of potentially cooperative genes. Consequently, the TIME cell monolayer on each microcarrier has to be monoclonal, having evolved from a single cell. To achieve a one-to-one cell-microcarrier ratio, we used a seeding ratio of one cell per two microcarriers. Starting from a single cell, TIME cells reached confluence in nine days, forming tightly sealed monolayers similar to cells grown from a larger initial number (Fig. 3b).

**Figure 3:**
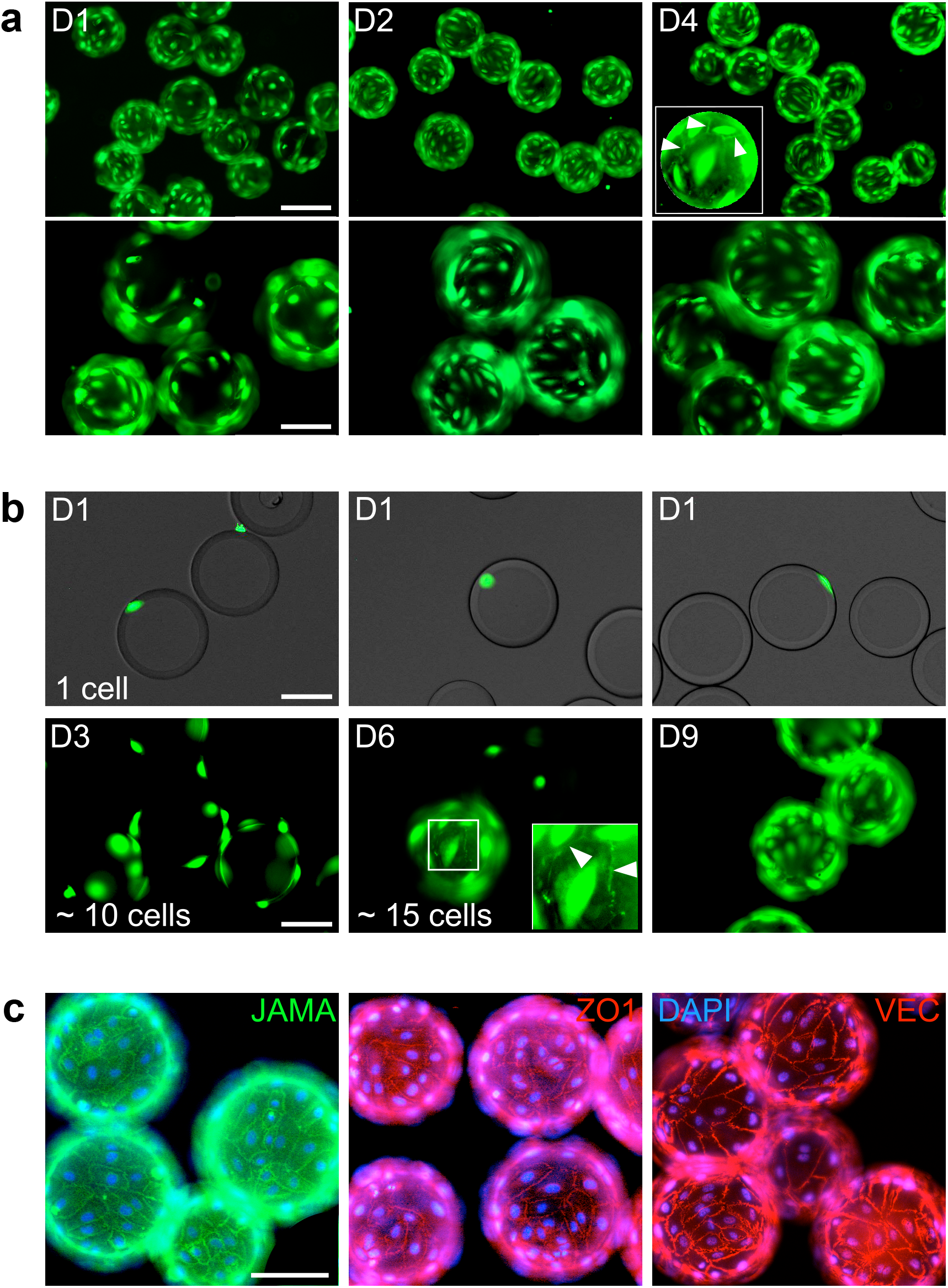
TIME cells form a clonal confluent cobble-stone monolayer on microcarriers. **a.** Images of calcein AM-stained TIME cells seeded on microcarriers at a density of 1.5×10^4^ cells/cm^2^ microcarrier surface area and grown for the indicated durations in spinner flasks. The cells reached confluence after 4 days and formed a tightly-sealed monolayer (see cell junctions marked by arrowheads in inset). Scale bars, 200 (top row) and 100 (bottom row) μm. **b.** When initiated from a single TIME cell per microcarrier, a confluent monolayer was formed in 9 days. The inset visualizes intercellular junctions marked by arrowheads. Scale bar, 100 μm. **c.** Microcarriers coated by confluent TIME monolayers immunolabeled with the indicated antibodies. Scale bar, 100 μm.

### TIME cells grown on microcarriers form intact tight and adherens junctions

Formation and maintenance of a cobblestone monolayer with integral intercellular junctions is essential for the repurposing of cell-coated microcarriers as individual permeability assays. Evidence for the establishment of such monolayers on the same type of microcarriers used here, including at the ultrastructural level, had been reported 30 years ago^35,42^. To find if junction-specific proteins are present at TIME cell intercellular junctions, we probed the cells for the tight junction single-pass transmembrane protein JAMA, and for the adaptor protein ZO1, which bind each other^43^. Both proteins were abundantly present at the borders of all the cells that were visible in the field of view, appearing as narrow undisrupted bands (Fig. 3c). The adherens junction marker vascular endothelial (VE)-cadherin was similarly abundant and patterned (ibid.). We conclude that the intercellular junctions formed by MAPs do not differ from junctions formed by the same (Fig. 2c) or other^44^ ECs on flat surfaces.

### Thrombin increases attachment of FNc to gelatin-coated surfaces

To assess the performance of FN_cf_ as a probe (Appendix SI, Fig. S1a), we tested if the extent of its binding to a gelatin-coated substrate decreases as EC confluence increases. The fluorescence signal of the probe attached to a glass coverslip harboring a confluent TIME cell monolayer was miniscule in comparison to the signal emanating from a sub-confluent monolayer (Appendix SI, Fig. S1b). Thrombin treatment of confluent TIME cell monolayers generated large gaps between adjacent cells (Appendix SI, Fig. S1c). Bare, gelatin-coated microcarriers bound FN_cf_ avidly, whereas microcarriers coated by a sub-confluent EC monolayer bound only a fraction of the latter (Appendix SI, Fig. S1d).

### Likelihood of false discovery is reduced by removal of non-confluent microcarriers

Though the majority of TIME cell monolayers reached confluence even when grown from a single cell per microcarrier, it is practically impossible to ensure that the surface of all microcarriers is fully coated by cells. Elimination of microcarriers that have residual exposed surface patches is essential because the FN_cf_ probe would bind to the exposed surface, causing misidentification of the cells as intransigent to the sgRNA they had been transduced with, and the overlooking of potentially relevant genes. To eliminate sub-confluent microcarriers from the population, we sorted MAPs that were incubated with FN_cf_ in the absence of an agonist. MAP extinction profiles were frequently uniform across the imaged face of the microcarrier but peaked at the edges of individual MAPs (Fig. 4a). The peaks corresponded probably to the gelatin and TIME cell layers. The initial gating of the MAP population excluded debris or MAP clusters (Fig. 4b). The residual MAP population was gated again to remove MAPs whose peak fluorescence intensity was at least two-fold higher than the intensity of the highest counts (ibid.), assuming this would eliminate most of the non-confluent MAPs. The removed fraction amounted to 12 percent of the MAP population. Consequently, the peak-height histogram of the residual MAPs was confined to a narrow range (Fig. 4c). Visual inspection of samples from the initial and residual populations confirmed the presence of exceptionally bright MAPs in the former (Fig. 4d), but not in the latter (Fig. 4e).

**Figure 4.**
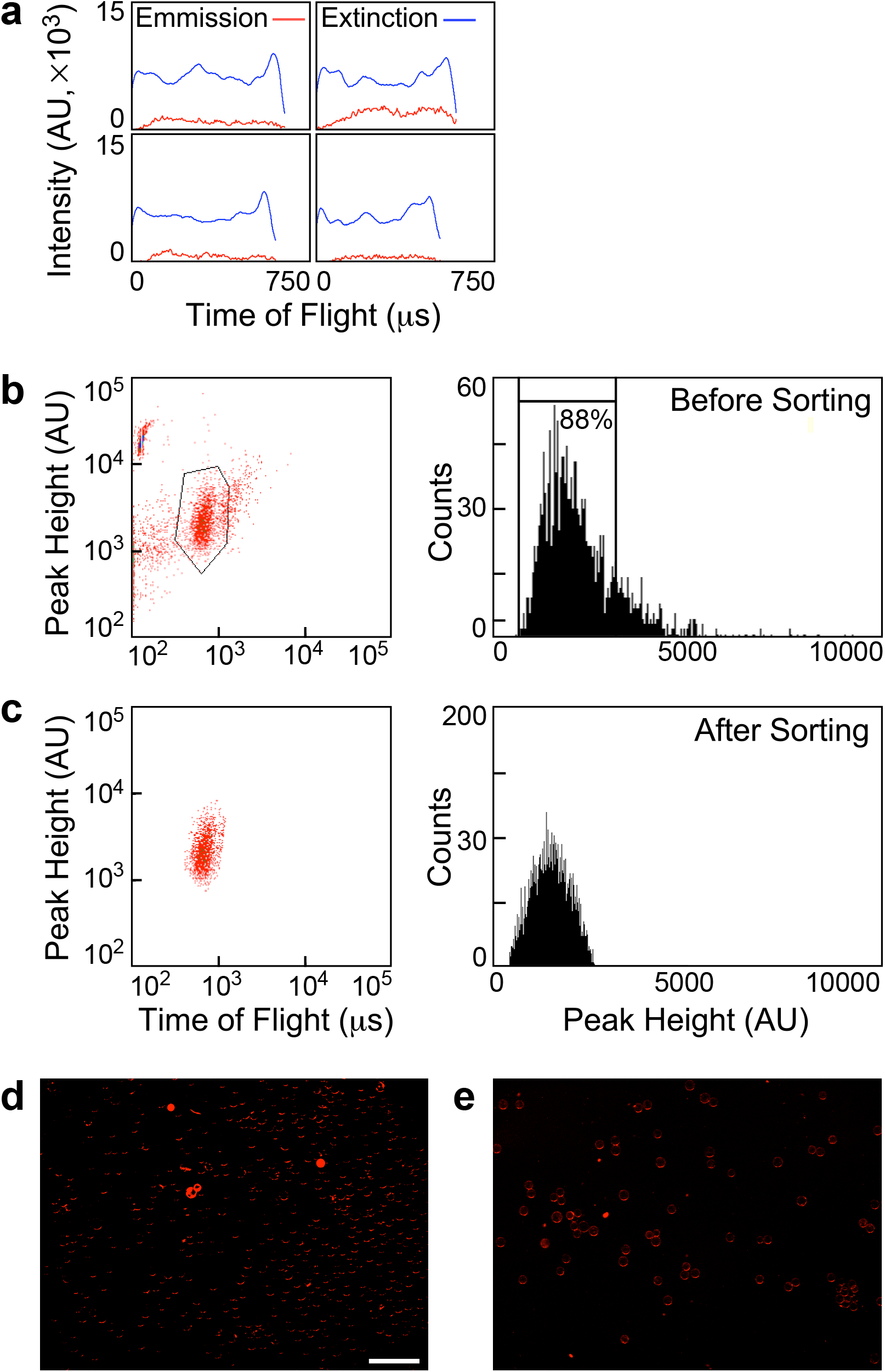
Binding of FN_cf_ removes non-confluent MAPs. **a.** Gallery of emission and extinction intensity profiles of individual MAPs. **b**. Left side: density plot of MAP extinction per time-of-flight of untreated MAP population that was incubated with FN_cf_ prior to sorting; right side: histogram of the MAP population shown on the density plot; the gate is placed at approximately 2-fold intensity of the median peak MAP fluorescence. **c.** Left side: density plot of the MAP population gated above; right side: histogram of the MAP population shown on the density plot. **d.** Image of a group of presorted MAPs showing the fluorescence of bound FN_cf_. **e.** A similar image of a group of sorted MAPs (scale bar, 1 mm).

### sgRNAs repress target gene expression level, protein abundance, and response to thrombin

To effectively test if the microcarrier-based permeability assay can identify genes that affect EC response to thrombin, we selected sgRNAs that target *F2R*, the gene that encodes the main thrombin receptor in ECs^45^, proteinase-activated receptor (PAR) 1, as a positive control. We chose *KDR*, the gene that encodes VEGF receptor 2, the main VEGF receptor in blood vessel ECs^46^, as a negative control. We included sgRNAs that target *GNAQ*, a gene encoding Gα_q_, a trimeric GTPase. Gα_q_ mobilizes calcium by activating phospholipase C-β, but, unlike Gα_12/13_, there is no consensus in regard to its ability to increase EC monolayer permeability^47,48^. We selected it, therefore, as an unknown. Before testing the effects of the selected sgRNAs on EC barrier function, we validated the presumed capacity of each of the five individual sgRNAs designed to target these genes^8^ by measuring how effectively they repressed the transcription of their target genes and, consequently, reduced the abundance of the proteins these genes encode. Using quantitative real-time polymerase chain reaction (qRT-PCR), we detected a large variability among the repressive effects within each set of five sgRNAs, which reached more than 13-fold in the case of *F2R*-targeting sgRNAs (Fig. 5a). The most effective sgRNA repressed *F2R* transcription to approximately 6 percent relative to its expression level in TIME cells treated by non-targeting sgRNAs (ibid.). Based on immunoblotting, the pattern of PAR1 abundance in each cell group expressing a single sgRNA fit closely the qRT-PCR expression pattern of *F2R* (which encodes PAR1) once the weight of each band was normalized relative to the weight of the corresponding loading-control β-actin band (Fig. S2a). The single sgRNA-specific expression levels of *GNAQ* and *KDR* fit well the abundances of Gα_q_ and VEGFR2 (Fig. 5b, 5c, Fig. 2b, 2c), confirming that two to three sgRNAs out of each set of five were capable of repressing their target gene by at least 60 percent.

**Figure 5.**
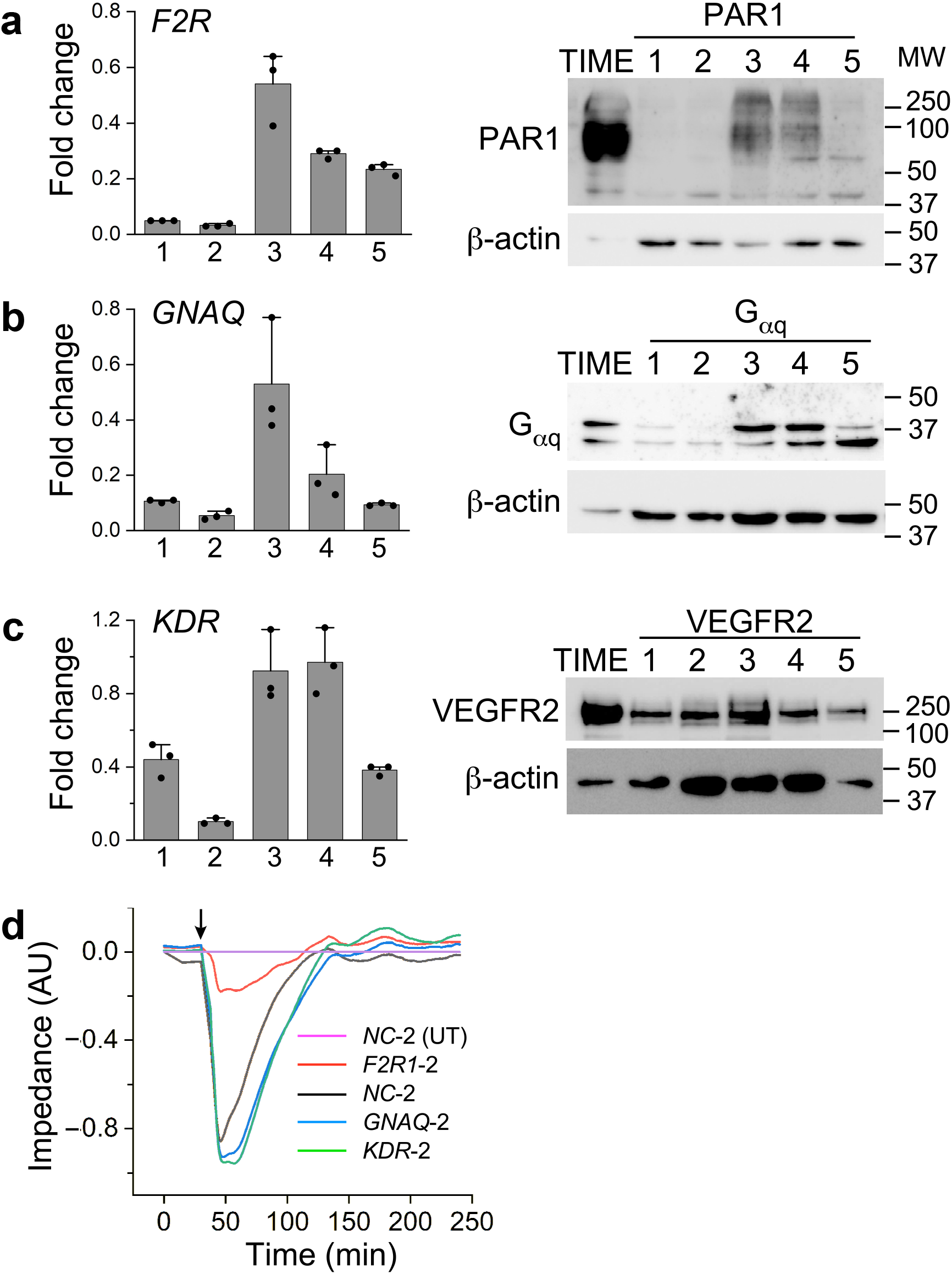
Highly effective sgRNAs reduce substantially the expression and abundance of the targeted gene and the encoded protein, respectively. **a**. Quantification of qRT-PCR measurements of the expression levels of *F2R* in TIME cells transduced by each one of the five CRISPRi library sgRNAs that target *F2R* (mean±SD, n=3). The immunoblot on the right shows PAR1 abundance in TIME cells transduced by the same sgRNAs. The first lane on the left was loaded with lysate of TIME cells that do not express sgRNAs. β-actin was used as a loading control for all immunoblots. **b** and **c**. The same measurements as above for the sgRNAs that target *GNAQ* or *KDR*, and the corresponding immunoblots. **d.** Impedance traces of TIME cells expressing each of the indicated sgRNAs. Thrombin was applied to all the cell groups at the time point marked by an arrow. The traces were normalized relative to the impedance of untreated (UT) cells expressing negative control (NC-2) sgRNA.

Efficient repression of *F2R* is expected to diminish the effect of thrombin on endothelial cell junctions, which typically lose their interconnections and retract, producing large intercellular gaps (Appendix SI, Fig. S1c). The effect of *GNAQ* repression, on the other hand, remains to be revealed in the planned permeability assays. To test these predictions, we measured the impedance of TIME cell populations expressing each of the most repressive sgRNAs of *F2R*, *GNAQ*, and *KDR*, grown in multi-well plates before and after the application of thrombin. The impedance of TIME cells expressing sgRNAs for *GNAQ*, *KDR*, and a validated negative-control (NC) sgRNA^8^ fell by close to one impedance unit (Fig. 5d). The impedance of TIME cells expressing *F2R* sgRNA, on the other hand, fell only by 0.2 units and returned to the basal level earlier than the other cell groups.

### Fluorescence-assisted sorting separates MAPs carrying thrombin-treated and untreated TIME cells

Execution of a genome-wide screen requires physical separation between the phenotypes that ensue during the functional assay. In the current study, these are thrombin-responsive and non-responsive MAPs. FN_cf_ would enable separation by fluorescence-assisted sorting. After thrombin treatment and before sorting, we fixed the microcarrier-attached TIME cells by EtOH, an optimal fixative for the preservation of nucleic acids^49^, to prevent changes in cell and junction morphology. Due to potential heterogeneity in cell response to thrombin, the light-intensity distribution of among the sorted MAPs was continuous over an interval of intensity values, rather than binary (Fig. 6a). To produce two distinct sorted populations, we defined two gates, one for the low-responding microcarriers, referred to as ‘negative’, and another for the high-responders that we refer to as ‘positive’ (ibid.). The fraction of untreated negative MAPs fell from approximately 40 percent to only 10 percent of the thrombin-treated microcarrier population because of the upward shift in MAP fluorescence intensity (Fig. 6b). Complementarily, the percentage of positive microcarriers increased from approximately 15 to 40 percent of the treated microcarrier population. These measurements demonstrate a robust and reproducible response to thrombin that can be readily resolved into positive and negative populations. A portion of approximately 45 percent of the total microcarrier population, encompassing the medium range of response, remained between the upper and lower gates. This sub-population is least likely to contain microcarriers carrying cells that express the most repressive sgRNAs. Sidestepping this population is not likely, therefore, to preclude the identification of genes that have major roles in mediating the disassembly of cell junctions in response to thrombin.

**Figure 6.**
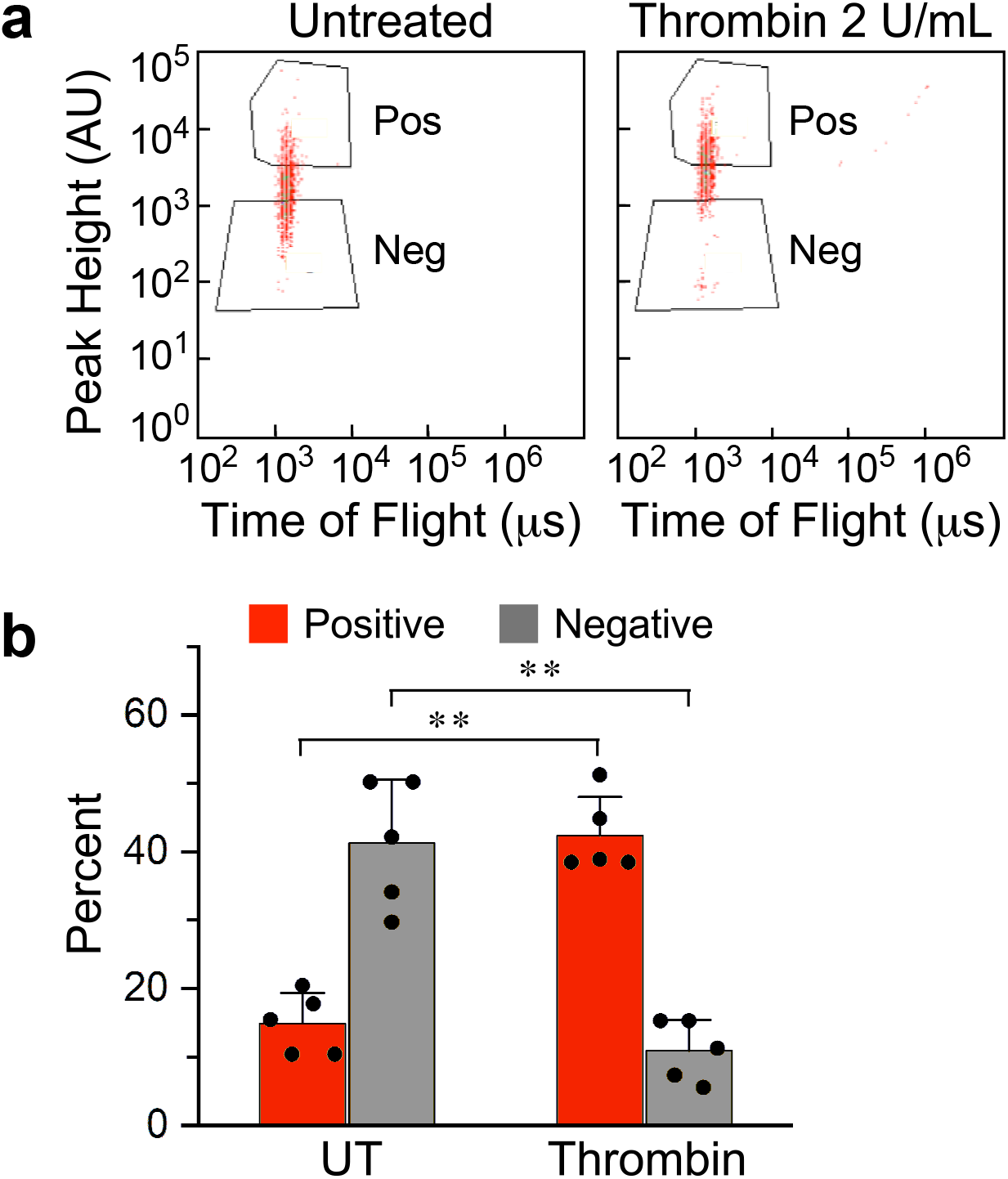
Dark and fluorescent microcarriers are over-abundant in untreated or thrombin-treated populations, respectively. **a**. Confluent TIME cells grown on microcarriers were either untreated or treated by 2 U/mL thrombin for 30 min followed by 5 min incubation with 1 μg/mL FN_cf_ for 5 min. The microcarriers were sorted according to the indicated gates into positive (Pos) and negative (Neg) subgroups. **b**. Histogram summarizing the negative and positive untreated or thrombin-treated populations of MAPs gated as shown in panel a (mean±SD, n=5, ** *P*<0.001).

### MAP sensitivity to *F2R* knockdown is similar to the sensitivity of impedance measurement

We chose impedance measurement as a benchmark for assay sensitivity because this technique is more sensitive than that of probe flux through cell monolayers cultured on permeable substrates^50^, the main alternative technique. We tested the sensitivity of impedance measurement as a detector of sgRNA-mediated knockdown by comparing the thrombin response of TIME cells transduced by the most effective *F2R*-targeting sgRNA, *F2R*-2, to cells transduced by a negative-control sgRNA. Whereas the impedance of cells transduced by the latter sgRNA fell by 2.6 units (denoted as I_NC_ in Fig. 7a), the impedance of TIME cells transduced by *F2R*-2 fell only by 0.75 unit (I_F2R_).

**Figure 7.**
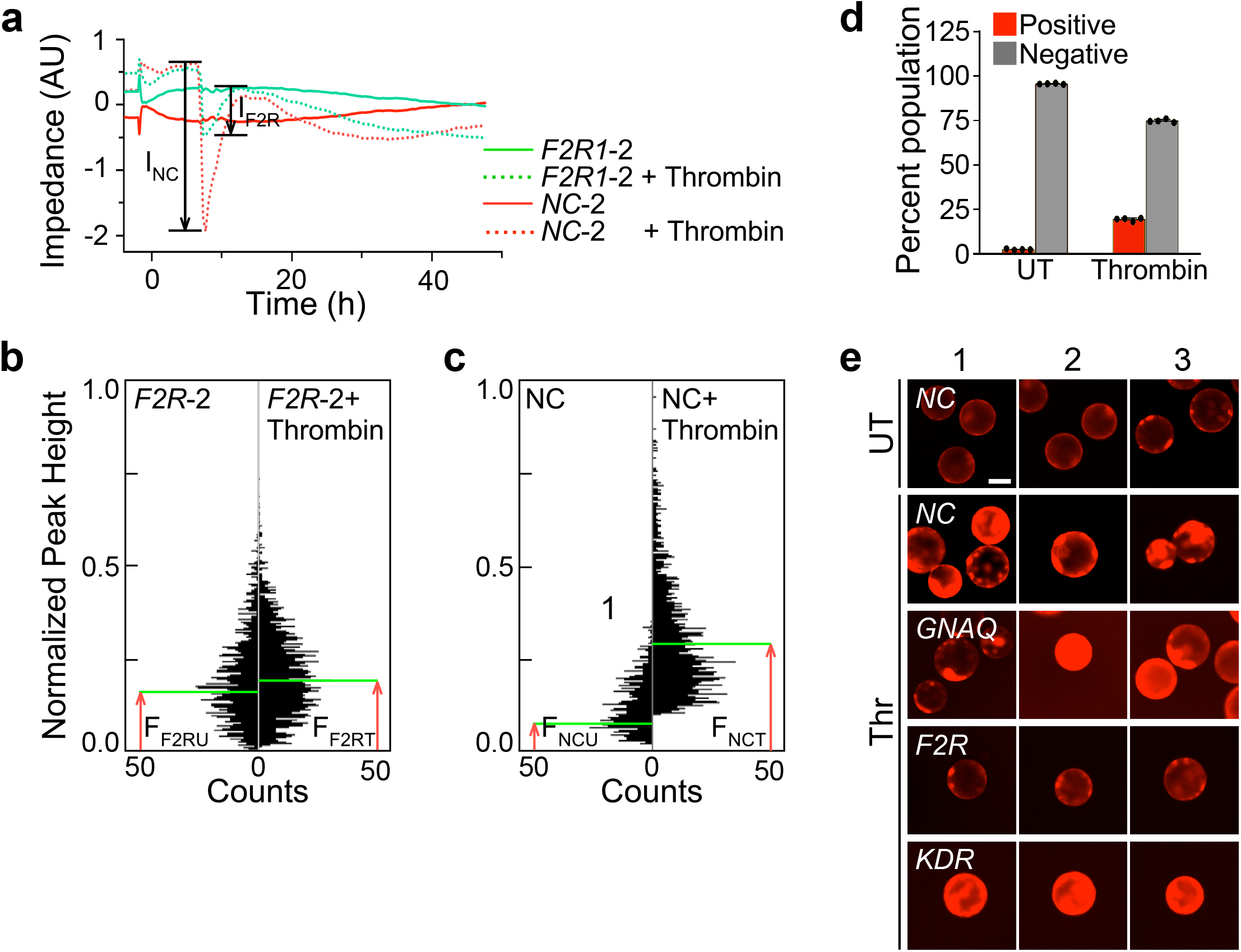
*F2R* sgRNA suppresses thrombin-induced hyperpermeability. **a**. Time course of impedance measurements of thrombin-treated TIME cells normalized relative to the impedance of untreated (UT) cells expressing negative control (NC-2) sgRNA. **b.** Gating of peak-height fluorescence density plot of MAPs carrying cells transduced by *F2R*-targeting sgRNA; righthand panel: display of the counts per peak height and gate of the same MAP population shown on the density plot. **c.** Counts per peak-height fluorescence of sorted MAPs that carry either untreated or thrombin-treated cells transduced by *F2R*-targeting sgRNA. **d.** Effects of thrombin treatment on the fluorescence intensity of MAPs carrying TIME cell transduced by the indicated sgRNAs. **e.** Triplicate fluorescence images of MAPs carrying cells transduced by the indicated sgRNAs under the indicated conditions. Scale bar, 100 μm.

As a first step in the test of MAP sensitivity to *F2R* knockdown, we repeated the procedure described above for the removal of particles that were not whole single MAPs and of non-confluent single MAPs (Fig. 4b). The mean peak fluorescence intensity of thrombin-treated MAPs carrying cells transduced by the most effective *F2R*-targeting sgRNA (denoted as F_F2RT_ in Fig. 7b) was higher by approximately 18 percent than that of the same type of untreated MAPs (F_F2RU_). On the other hand, MAPs carrying cells transduced by a mixture of three negative control sgRNAs responded to thrombin by a four-fold increase in mean fluorescence intensity compared to an untreated group (compare F_NCT_ to F_NCU_ in Fig. 7c). Because impedance measurement sensitivity is a dimensionless variable given by the ratio between I_NC_ and I_F2R_ (Fig. 7a), we defined MAP sensitivity as the ratio between the response of cells which express a mixture of three negative control sgRNAs and cells that express *F2R*-2, i.e. the ratio between F_NCT_–F_NCU_ (Fig. 7b) and F_F2RU_–F_F2RT_ (Fig. 7c). The respective sensitivities were 3.5 and 7.6, indicating that MAP sensitivity was comparable or better than the sensitivity of impedance measurement for detecting the repressive effect of the *F2R*-2 targeting sgRNA.

### *F2R* sgRNA suppresses thrombin-induced hyperpermeability

All the sgRNAs tested above repress, albeit to varying degrees, the expression of their target genes. Therefore, we gauged the combined effect of these sgRNAs on MAP permeability by comparing the fraction of MAPs sorted from a mixed population carrying cells that express on each MAP one of the 15 sgRNAs employed in the study (five each for *F2R*, *GNAQ*, and *KDR*) to the fraction of MAPs carrying cells that express negative control sgRNAs. The sorting of the first group (Fig. 7d) showed that sgRNA expression almost abolished the population of untreated positive MAPs and reversed the distribution of thrombin-treated MAPs shown in Fig. 6b, i.e. reduced the number of positive MAPs while increasing the number of negative MAPs. The former was more than halved, and the latter increased by close to seven-fold. These large effects demonstrate the repressive efficacy of each gene-specific group of five sgRNAs. To visualize the individual effect of the most repressive *F2R*, *GNAQ* and *KDR*-targeted sgRNAs on MAP permeability in comparison to three validated NC sgRNAs^8^, we imaged the fluorescence of thrombin-treated MAPs expressing each of these sgRNAs. Cells expressing either *GNAQ*, or *KDR*-targeted sgRNAs responded to thrombin similarly to negative-control sgRNA-expressing MAPs (Fig. 7e). In contrast, the response of MAPs carrying cells that expressed the most effective *F2R* sgRNA was similar to that of MAPs carrying untreated cells that expressed one of the negative-control sgRNAs (top row of images in Fig. 7e). These results are consistent with the other functional effects of the sgRNAs shown above. The persistence of low fluorescence intensity emitted by MAPs carrying cells that express a *F2R* sgRNA indicates that thrombin treatment did not cause an artifactual signal resulting from cell detachment from the microcarrier surface. Furthermore, an assay with MAPs that express a repressive sgRNA library requires sorting of microcarriers that emit weak light signals. Therefore, artifactually bright microcarriers would not cause false gene identification.

## DISCUSSION

Permeability is not amenable to existing functional assays, because it is inherently a collective cell function. The new assay repurposes a pre-existing device – microcarriers that had been developed over fifty years ago to increase the surface available for the growth of adherent cells^51^. Microcarriers had already been used, in fact, to measure EC monolayer permeability, albeit in bulk format^35^. In contrast, the new assay deploys each cell-coated microcarrier as a single sample, thus taking advantage of its small diameter to pack as many as 6000 microcarriers per mL of medium. At this sample concentration, 166 mL of growth medium would contain 10^6^ microcarriers. As estimated above, this is a sufficiently large number of samples for performing a genome-wide screen with this functional assay.

Screens based on cell growth can measure a continuum of quantitative phenotypes by counting the numbers of cells that express sgRNAs of interest^52^. These measurements can be used for further analysis of the targets identified in the first screen, such as construction of genetic interaction (GI) maps^53^. Though the phenotypes of the MAPs (i.e., their fluorescence intensities), also vary continuously within a certain range, the sorting converts that range into an either ‘positive’ or ‘negative’ binary output, precluding the attribution of a continuous range of quantitative phenotypes to each sample, and of further analysis. This drawback can be overcome, however, by performing a set of assays where the position of the gate is changed incrementally so that a larger range of phenotypes is sampled. Replicating the assay is straightforward because it can be run in as little as 30 min using a volume of less than 200 mL microcarriers in growth medium.

In addition to compactness, the format of the assay facilitates physical separation between samples that are either responsive or refractory to the stimulus, by using the phenotype as a sorting input. The separation is crucial for the identification of the genes that produce the measured phenotype. Counting the number of cells on the microcarriers, and determining if the cell monolayer reached confluence are straightforward because the cells on the microcarrier surface can be readily visualized by a dye and by other imaging modalities^35,42^. The visibility enables growth of a clonal cell population from a single cell per MAP and makes it possible to verify that the cell monolayer coating the MAPs is confluent, preventing spurious signal as a result of probe binding to the exposed surface of the microcarrier. The duration of the assay, i.e. the time necessary for eliciting a measurable phenotype, depends on the nature of the agonist, but it is not likely to allow sufficient time for the emergence of multiple phenotypes, such as change in cell density. This narrows the scope of the assay to its intended purpose of measuring intercellular permeability and prevents interference from other potential effects of the agonists which are often pleiotropic, such as VEGF.

As is standard practice in pioneering proof-of-principle studies, we favored permissive experimental conditions. Accordingly, we chose an agonist that induces relatively large gaps that did not strain the detection limits of our probe. Admittedly, other agonists produce subtler remodeling of intercellular junctions, requiring higher sensitivity for distinguishing between the signals emitted by monolayers of quiescent and stimulated ECs. As long as the agonist induces sufficiently large openings between adjacent cells to permit binding of the probe to the microcarrier surface, our approach should be feasible as long as the binding affinity between the probe and the surface is sufficiently high. VEGF, an obvious agonist of interest, generates gaps on the order of several micrometers^54^. Its signaling pathway should be amenable, therefore, to MAP screening. The MAP configuration limits to some extent the range of testable conditions because it does not simulate abluminally-acting agonists, including, in some cases, VEGFR2-downstream effects^55^. Genes that are required for cell proliferation and for adhesion to gelatin will not be represented in the screened cell population because cells that express sgRNAs which target these genes will be lost during cell growth on microcarriers. It is unlikely, however, that such genes would be directly involved in regulating intercellular junction integrity.

## MATERIALS AND METHODS

### Endothelial cell culture

TIME cells (American Type Cell Culture) were grown at 37°C in a humidified 5 percent CO_2_ incubator in Vascular Cell Basal Medium supplemented with Microvascular Endothelial growth Kit-VEGF (American Type Cell Culture) and 12.5 μg/mL blasticidin (ThermoFisher Scientific).

### Immunofluorescence

TIME cells grown to confluence on gelatin-coated round 12-mm glass coverslips (GG-12-Gelatin, Neuvitro) were treated with 2 U/mL thrombin (human α-thrombin, Enzyme Research Laboratories) for 30 min at 37°C, and 5 percent CO_2_, or with PBS as a negative control. After washing in ice-cold PBS, fixation in 4 percent paraformaldehyde/PBS for 30 min at 23°C, and permeabilization in 0.1 percent Triton X-100 (Sigma-Aldrich) in PBS for 5 min, cells were incubated in a blocking solution of 5 percent BSA (Sigma-Aldrich) in PBS for 1 h. Immunolabeling by primary antibodies diluted according to the manufacturer’s instructions in 0.1 percent Triton X-100 and1 percent BSA in PBS overnight at 4 °C was followed by a 3×PBS wash and incubation with 2.5 μg/mL fluorescently-conjugated secondary antibodies in PBS for 30 min at 23°C. After a 3×PBS wash, coverslips were mounted on glass slides (ProLong Gold Antifade Mountant, ThermoFisher Scientific). Images of TIME cells grown either as a two-dimensional culture or on microcarriers were acquired by epi-fluorescence (EVOS FL Auto Cell Imaging System, ThermoFisher Scientific) or by laser-scanning confocal microscopy (Nikon A1R+).

### Immunoblotting

Cells were scraped off plates, extracted in RIPA lysis buffer (25 mM Tris-HCl pH 7.6, 150mM NaCl, 1 percent NP-40, 1 percent sodium deoxycholate, 0.1 percent SDS; ThermoFisher Scientific) supplemented with phosphatase (Fisher Scientific) and protease (Sigma-Aldrich) inhibitor cocktails, and clarified by centrifugation (20,000 g, 10 min). Total protein concentration was determined by BCA protein assay (ThermoFisher Scientific). Equal amounts of total protein were separated by 10 percent SDS-PAGE and transferred to a PVDF membrane. Membranes were blocked by 5 percent casein in TBS with 0.1 percent Tween 20 (Sigma-Aldrich), probed with the indicated primary antibodies, and incubated with HRP-conjugated secondary antibodies for 1 h at 23°C. Protein-antibody complexes were visualized by chemiluminescence (SuperSignal™ West Pico PLUS, ThermoFisher Scientific) and imaged by a charge-coupled device camera (Chemidoc Imaging System, BioRad). Band densitometry was done with the Gels submenu of ImageJ 2.0.0.

### Impedance measurement

Impedance was measured on xCELLigence Real-Time Cell Analyzer (ACEA Biosciences) according to the manufacturer’s instructions. Briefly, 30000 TIME cells were seeded in 200 μL growth medium per well of an electronic 16-well plate (E-Plate View 16, ACEA Biosciences) for 30 min at 23°C to allow cell sedimentation. The cells were grown to confluence at 37°C, 5 percent CO_2_. On the day of the experiment, cells were treated by thrombin or by the same volume of carrier (PBS), and placed in the analyzer for continuous recording of impedance. The relative impedance output of the instrument is expressed as (Z_i_-Z_0_)/15 Ω, where Z_i_ is the impedance at the i^th^ time point, and Z_0_ is the initial impedance before agonist addition.

### Macromolecular permeability

TIME cells seeded at 1×10^5^ per filter insert (0.4 μm pores, 6.4 mm diameter, polyethylene terephthalate membrane, Transwell®, Corning) were grown for 72 h in 24-well plates (Transwell®, Corning) filled with 800 μL growth medium per well. The cells in the inserts were supplemented at the same time by either 2 U/mL thrombin or carrier, and by 1 μg/mL fluorescein isothiocyanate (FITC)-dextran (2000 kDa, Sigma-Aldrich) was added in the upper chamber. Ten μL samples were removed after 0, 5, 15, 30, 60, 90 and 120 min from the lower compartment, diluted in 90 μL of PBS, transferred in a 96 well black plate, clear bottom. Fluorescence in each well was measured with excitation at 485 nm and readout at 535 nm (VICTOR2, PerkinElmer). The results are expressed as light intensity counts.

### TIME cell culture on microcarriers in spinner flasks

Cytodex-3 microcarriers (GE Health Sciences) were hydrated in PBS at 50 mL/g for 3 h at 23°C, washed twice in a 30 mL/g PBS, and autoclaved (121°C, 20 min). A quantity of 92 mg cytodex-3 microcarriers were readied for cell growth by incubation overnight at 37°C, 5 percent CO_2_, and stirred at 35 rpm (Troemner) in 50 mL growth medium in siliconized (Sigmacote, Sigma-Aldrich) spinner flasks (Bellco Glass, model 1965-95010) equipped with a glass-ball plunger. TIME cells grown to 80 percent confluence in T75 flasks were briefly detached with 0.05 percent trypsin, 0.02 percent EDTA (American Type Cell Culture). The cells were then resuspended in growth medium supplemented with 15 percent FBS and 4.5 g/L glucose. Microcarriers were inoculated with 6×10^6^ cells in 50 mL same growth medium to achieve a density of 24000 cells/cm^2^. The microcarrier culture was incubated at 37°C, 5 percent CO_2_ and stirred intermittently at 35 rpm for 1 min once every 15 min for a total of 1 hour, then stirred continuously at the same speed. Medium was replenished once every two days. Cells reached confluence in 3-5 days. In order to seed a single cell per microcarrier, the above amount of microcarriers was inoculated with 1.5×10^5^ cells to achieve a density of 600 cells/cm^2^. To monitor cell growth on microcarriers, a small microcarrier sample was removed every day, treated with 2 μM of Calcein AM (ThermoFisher Scientific), and visualized by epifluorescence microscopy. Under these conditions, cells reached confluence in 9 days.

### Detection of barrier disruption by fluorophore-conjugated fragment of fibronectin

To detect disruption of the confluent TIME cell monolayer on microcarriers, we used a 45 kDa FN_c_ (Sigma-Aldrich) purified from human plasma (Sigma-Aldrich) conjugated to Dylight 550 with a labeling kit (ThermoFisher Scientific). The cells were grown to confluence on gelatin-coated round 12-mm diameter glass coverslips (GG-12-Gelatin, Neuvitro). On the day of the permeability assay, the cells were treated with thrombin for 30 min at 2 U/mL and with 1 μg/mL Dylight 550-conjugated FN_cf_ for 5 min. The cells were then washed with ice cold PBS, fixed in 4 percent paraformaldehyde/PBS for 30 min, and washed again with PBS. The coverslips were mounted on glass slides with Prolong Gold (Invitrogen), a DAPI-containing medium. Permeability was characterized by quantifying dye fluorescence (DyLight 550, Ex/Em 562/576 nm, ThermoFisher Scientific) in epifluorescence images (EVOS FL, ThermoFisher Scientific).

### Permeability in response to thrombin on cytodex-3 microcarriers

TIME cells grown to confluence on microcarriers were incubated with 2 U/mL of Thrombin for 25 min in spinner flasks spun at 35 rpm at 37°C, 5 percent CO_2_. The medium was then supplemented with 1 μg/mL FN_cf_ and stirred for 5 min. MAPs were transferred to a glass-bottom plate for visualization and quantification of the Dylight 550 fluorescence (EVOS FL).

### Microcarrier sorting

MAPs treated as required were fixed in 50 percent EtOH/PBS and 1 percent FBS for 10 min at 23°C, washed twice in PBS for 10 min at 4°C, and transferred to PBS with 1 percent FBS. MAPs were sorted at 561 nm on a Biosorter instrument (Union Biometrica) equipped with a FP-1000 nozzle.

### Expression of *dCas9-KRAB* and repressive sgRNAs in TIME cells

Lentiviri were produced and infected into TIME cells as described^56^. The cells were transduced first by lentivirus expressing dCas9 fused to the KRAB repressor module^9^ (pHR-SFFV-dCas9-BFP-KRAB, Addgene). Lentivirus-expressing cells were sorted (FACSAria II, BD Biosciences) using blue fluorescent protein (BFP) as marker. Oligonucleotides encoding five optimized sgRNAs targeting the transcription start sites of *F2R*, *GNAQ*, and *KDR*, and NC sgRNAs were synthesized based on sequences from the CRISPRi V2 library designed by Horlbeck et al.^8^. Each sgRNA was subcloned into the backbone plasmid of the CRISPRi V2 library (pU6-sgRNA EF1Alpha-puro-T2A-BFP, Addgene) between the BstXI and BlpI restriction sites. Groups of *dCas9-KRAB*-expressing TIME cells were each infected separately by lentiviri expressing a single sgRNA and selected by puromycin resistance conferred by the sgRNA-expressing plasmid. sgRNA sequences are provided in Appendix SI, Table S1.

### Statistics

The significance of the difference between means was determined by two-tailed Student’s *t*-test. The null hypothesis was considered untrue if the probability satisfied the condition *P*≤0.05.

## Non-standard Abbreviations

BFP: blue fluorescent protein
CRISPRi: inhibitory clustered regularly interspaced short palindromic repeats sgRNA library
dCas9: deactivated CRISPR-associated protein 9
DAPI: 4′,6-diamidino-2-phenylindole
EC: endothelial cell
FITC: fluorescein isothiocyanate
FN_c_: collagen-binding fragment of fibronectin
FN_cf_: fluorophore-conjugated FN_c_
KRAB: Krüppel-associated box (KRAB)
MAP: microcarrier assay of permeability
PAR: proteinase-activated receptor
qRT-PCR: quantitative real-time polymerase chain reaction
sgRNA: single guide RNA
TIME: telomerase-immortalized microvascular endothelial
UT: untreated
VE: vascular endothelial
VEGF: vascular endothelial growth factor
VP64: tetrameric VP16 transactivation domain

## DECLARATIONS

### Ethics approval and consent to participate

No animals and no humans participated in this study.

### Consent for publication

All authors consented for the publication of this study.

### Availability of data and material

All data and materials generated in this study will be made available to the scientific community.

### Competing Interests

The authors have no competing financial or other interests.

### Funding

This study was supported by the National Heart Lung and Blood Institute (R01 grant HL119984 to AH and R01 grant HL128234 to LE) and by the National Cancer Institute (P30 grant CA56036 to PF).

### Author Contributions

CS designed and performed experiments, analyzed the results, and wrote the manuscript; JY and XK performed experiments, analyzed the results, and prepared reagents; RK performed experiments, LE supervised the preparation of reagents and designed experiments; PF supervised the preparation of reagents and DNA sequencing, and analyzed data; EL designed experiments, analyzed data, and wrote the manuscript; AH conceived the study, coordinated its performance, designed experiments, analyzed the results, and wrote the manuscript.

## Acknowledgments

We thank Dr. Florin Tuluc, Flow Cytometry Core Laboratory of the Children’s Hospital of Philadelphia, for assistance in sorting; Drs. Luke Gilbert and Max Horlbeck, and Ryan Pak, for assistance in the use of the CRISPR/dCas9 library developed in the lab of Dr. Jonathan Weissman, University of California, San Francisco. We thank Dr. Jonathan Weissman for depositing the CRISPRi library and dCas9-KRAB plasmid at Addgene.

## SUPPLEMENTAL MATERIAL

### TABLE LEGEND

**Table S1.**
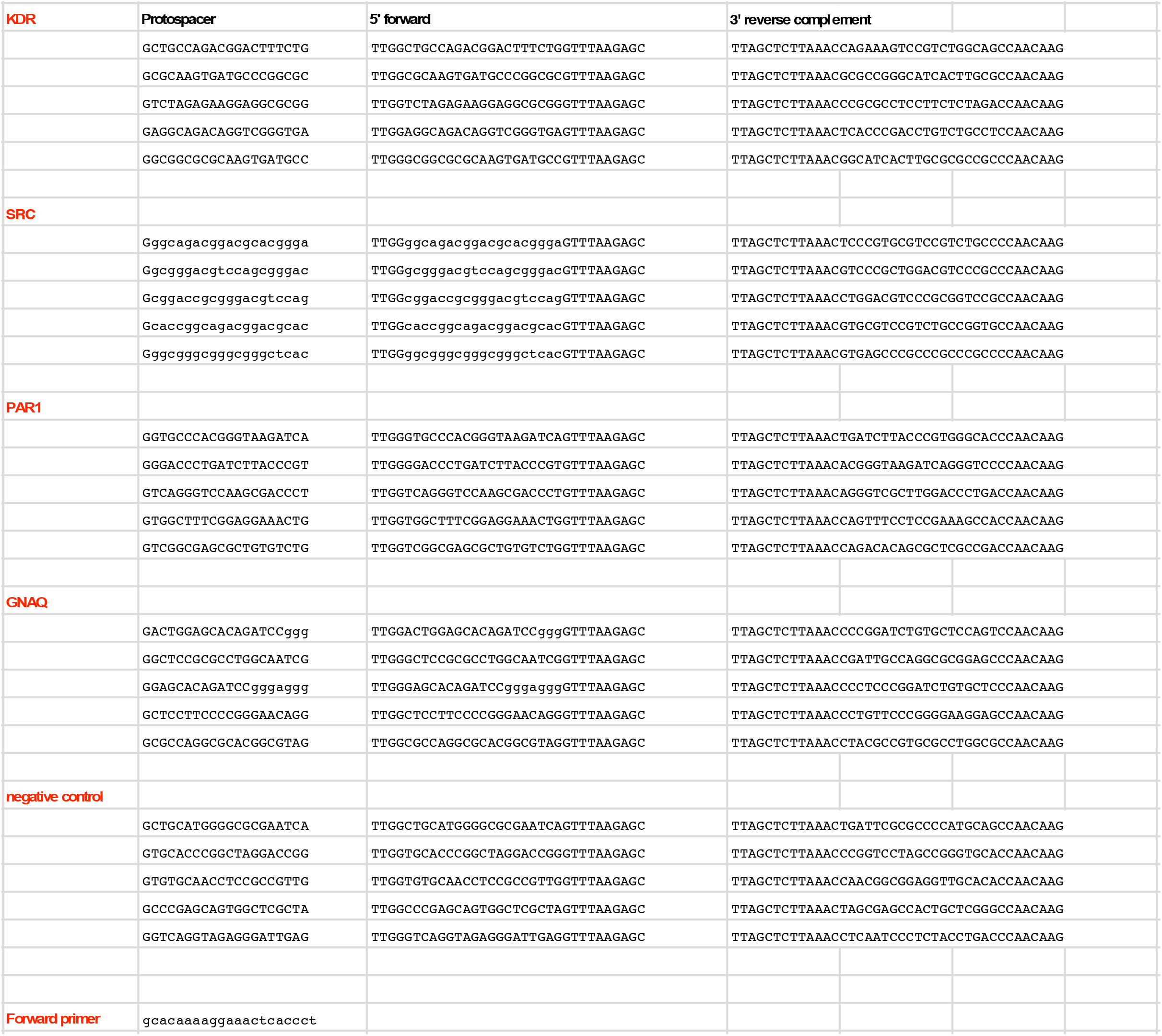
Sequences of the sgRNAs from the CRISPRi V2 library used in the study, the sgRNA-specific reverse primers used in qRT-PCR experiments, and the forward primer based on the sequence of the backbone plasmid of the sgRNA library, pU6-sgRNA EF1Alpha-puro-T2A-BFP.

### FIGURE LEGENDS

**Figure S1:**
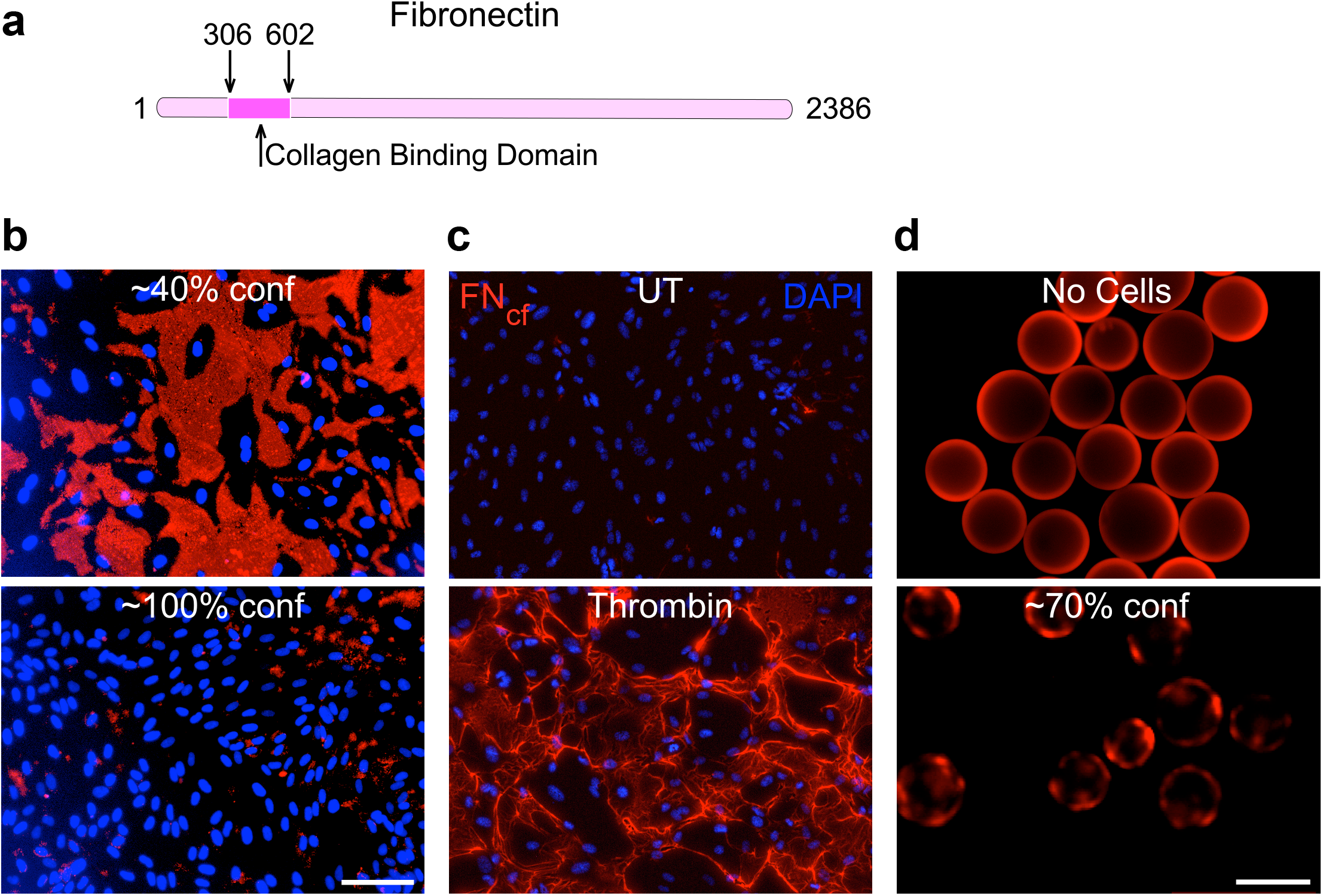
FN_cf_ identifies gaps between thrombin-treated confluent TIME cells. **a.** Scheme of human fibronectin showing the location of the collagen-binding domain. Numbers indicate amino acid positions. **b**. TIME cells grown on gelatin-coated glass coverslip until reaching the indicated confluences were incubated with 1 μg/mL FN_cf_ (red) for 5 min. Nuclei were stained by DAPI (blue). Scale bar, 100 μm. **c.** Confluent TIME cells grown as described above were either untreated (UT) or treated by 2 U/mL thrombin (Thr) for 30 min, followed by incubation with 1 μg/mL FN_cf_ for 5 m. **d.** Microcarriers devoid of cells (top) or carrying TIME cells at the indicated confluence (bottom) were incubated with 1 μg/mL FN_cf_ for 30 minutes. Scale bar, 200 μm.

**Figure S2:**
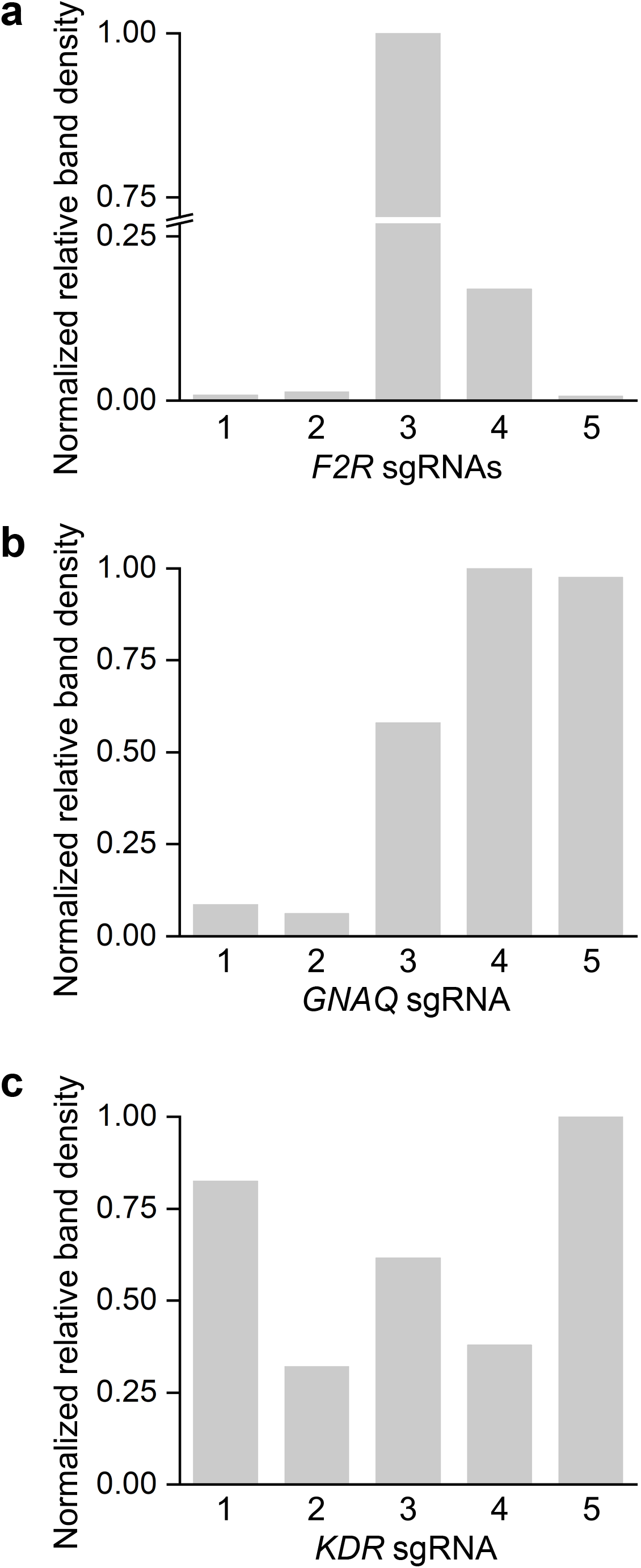
The pattern of sgRNA-induced decrease in protein abundance is similar to the pattern of the decrease in gene expression. **a.** Densitometric quantification of PAR1 abundance in cells expressing each of the 5 F2R-targeting sgRNAs shown in Fig. 5a, after normalization by the weights of the corresponding β-actin loading-control bands. **b** and **c.** Densitometric quantification similar to PAR1 of Gα_q_ and VEGFR2 abundances, respectively.

**Figure S3.**
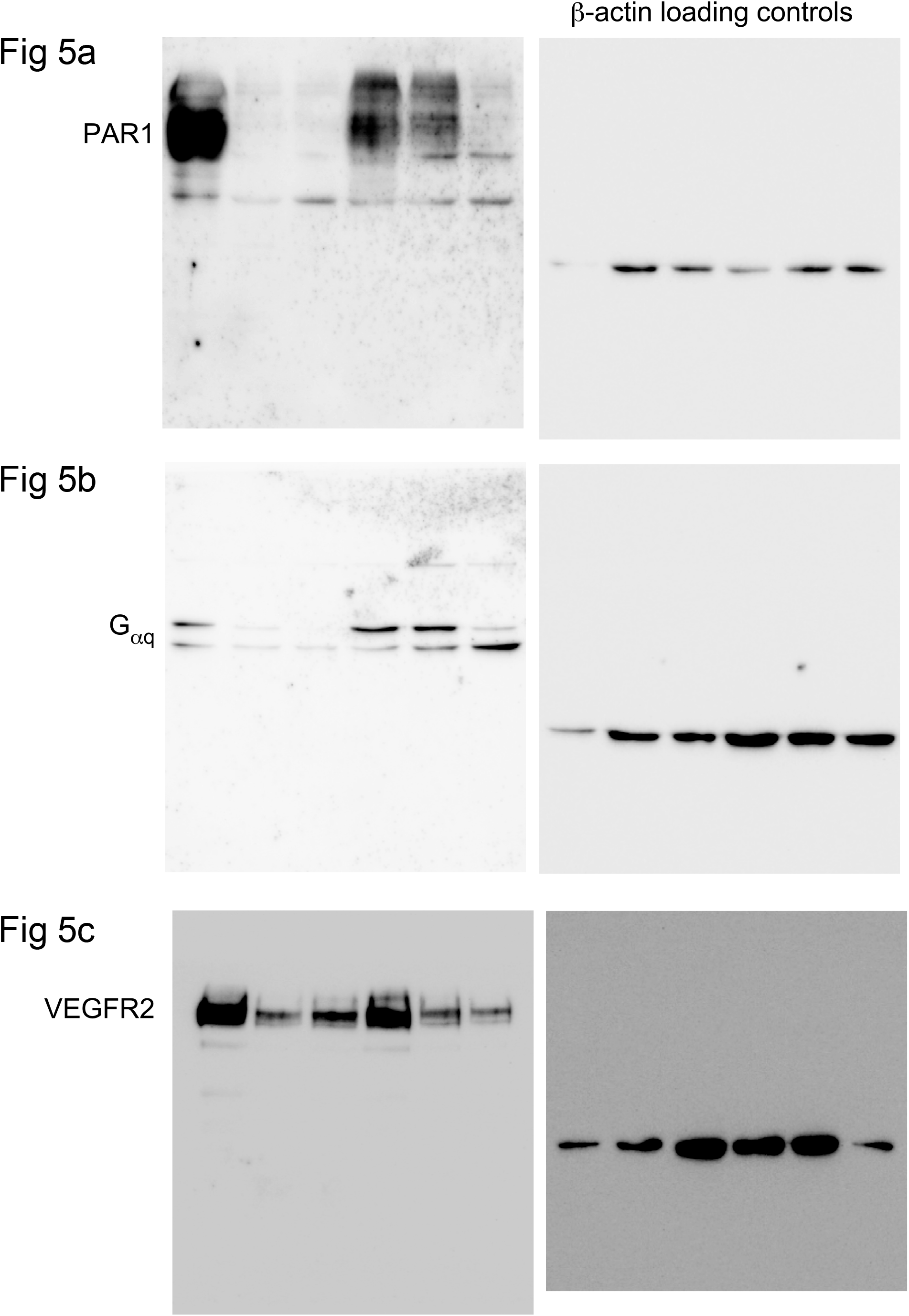
Full-length immunoblots of the bands shown in Fig. 5. The panel to which each immunoblot corresponds is indicated.

## Notes

#### Summary of Updates

Revised text and the figures, and and added supplementary data as Fig S2 in response to reviewers' comments.

